# H2AX promotes replication fork degradation and chemosensitivity in BRCA-deficient tumours

**DOI:** 10.1101/2023.05.22.541781

**Authors:** Diego Dibitetto, Martin Liptay, Francesca Vivalda, Ewa Gogola, Hulya Dogan, Martín G. Fernández, Alexandra Duarte, Jonas A. Schmid, Stephen T. Durant, Josep V. Forment, Ismar Klebic, Myriam Siffert, Roebi de Bruijn, Arne N. Kousholt, Nicole A. Marti, Martina Dettwiler, Claus S. Sørensen, Massimo Lopes, Alessandro A. Sartori, Jos Jonkers, Sven Rottenberg

**Author notes:** These authors contributed equally (Diego Dibitetto, Martin Liptay).

## Abstract

Histone H2AX plays a key role in DNA damage signalling in the surrounding regions of DNA double-strand breaks (DSBs)^1,2^. In response to DNA damage, H2AX becomes phosphorylated on serine residue 139 (known as γH2AX), resulting in the recruitment of the DNA repair effectors 53BP1 and BRCA1^3–6^. Here, by studying resistance to poly(ADP-ribose) polymerase (PARP) inhibitors in BRCA1/2-deficient mammary tumours^7,8^, we identify a novel function for γH2AX in orchestrating drug-induced replication fork degradation. Mechanistically, γH2AX-dependent replication fork degradation is elicited by the inhibition of CtIP-mediated fork protection. As a result, H2AX loss restores replication fork stability and increases chemoresistance in BRCA1/2-deficient tumour cells without restoring homology-directed DNA repair, as highlighted by the lack of DNA damage-induced RAD51 foci. Furthermore, in the attempt to discover acquired genetic vulnerabilities, we find that ATM inhibition overcomes PARP inhibitor (PARPi) resistance in H2AX-deficient tumours by interfering with CtIP-mediated fork protection of stalled forks. In summary, our results demonstrate a novel role for H2AX in replication fork biology in BRCA-deficient tumours and establish a function of H2AX separable from its classical role in DNA damage signalling and DSB repair.

---

To identify new genes involved in the cellular response to PARPi in BRCA2;p53-deficient tumours, we performed a genome-wide CRISPR-Cas9 screen in our KB2P3.4 (*K14cre*;*Trp53*^*-/-*^*;Brca2*^*-/-*^) mouse mammary tumour cell line^7^. Cells were transduced with the mouse sgRNA GeCKO_V2 library targeting 20,628 genes^9^, and then treated with a nearly lethal dose of 200nM of the AZD2461 PARPi for 3 weeks (Fig. 1a). At the end of the treatment, extracted genomic DNA from surviving cells was subjected to NGS and analyzed with the MAGeCK MLE algorithm^10^ (Fig. 1a). To increase the confidence of our screening data, we crossed the hits of this screen with the results obtained from four other genetic screens: two screens for PARPi resistance carried with a targeted DDR shRNA library in KB2P3.4 cells treated with AZD2461 or olaparib^11^; a genome-wide CRISPR-Cas9 screen performed in human RPE1-h*TERT TP53*^*-/-*^*;BRCA1*^*-/-*^ cells selected with olaparib^12^; a genome-wide CRISPR-Cas9 screen recently performed in our lab in KB2P1.21 cells (*K14cre*;*Trp53*^*-/-*^*;Brca2*^*-/-*^) treated with a lethal dose of cisplatin (*Widmer et al. manuscript in preparation*) (Fig. 1b). This data processing allowed us to identify general chemoresistance mechanisms independent of: (1) the specific PARPi, (2) BRCA1 or BRCA2 deficiency, (3) the type of screen (shRNA- or CRISPR-based), (4) the species (mouse or human), or (5) the anti-cancer agent used (PARPi or cisplatin). Our analysis revealed that sgRNA/shRNA against the histone *H2afx*/*H2AFX* were greatly enriched in all the analyzed screens and scored among the top identified hits (Fig. 1b, c). Conversely, sgRNA against 53BP1, a known key modulator of chemoresistance in BRCA1-deficient tumours, were enriched specifically in the RPE1*-*h*TERT TP53*^*-/-*^*;BRCA1*^*-/-*^ screen but not in the screens performed in BRCA2-deficient cells (Fig. 1c). These results suggest that H2AX loss is associated with PARPi resistance through a mechanism different than 53BP1 inactivation which only occurs in BRCA1-but not in BRCA2-deficient cells^13^.

**Fig. 1.**
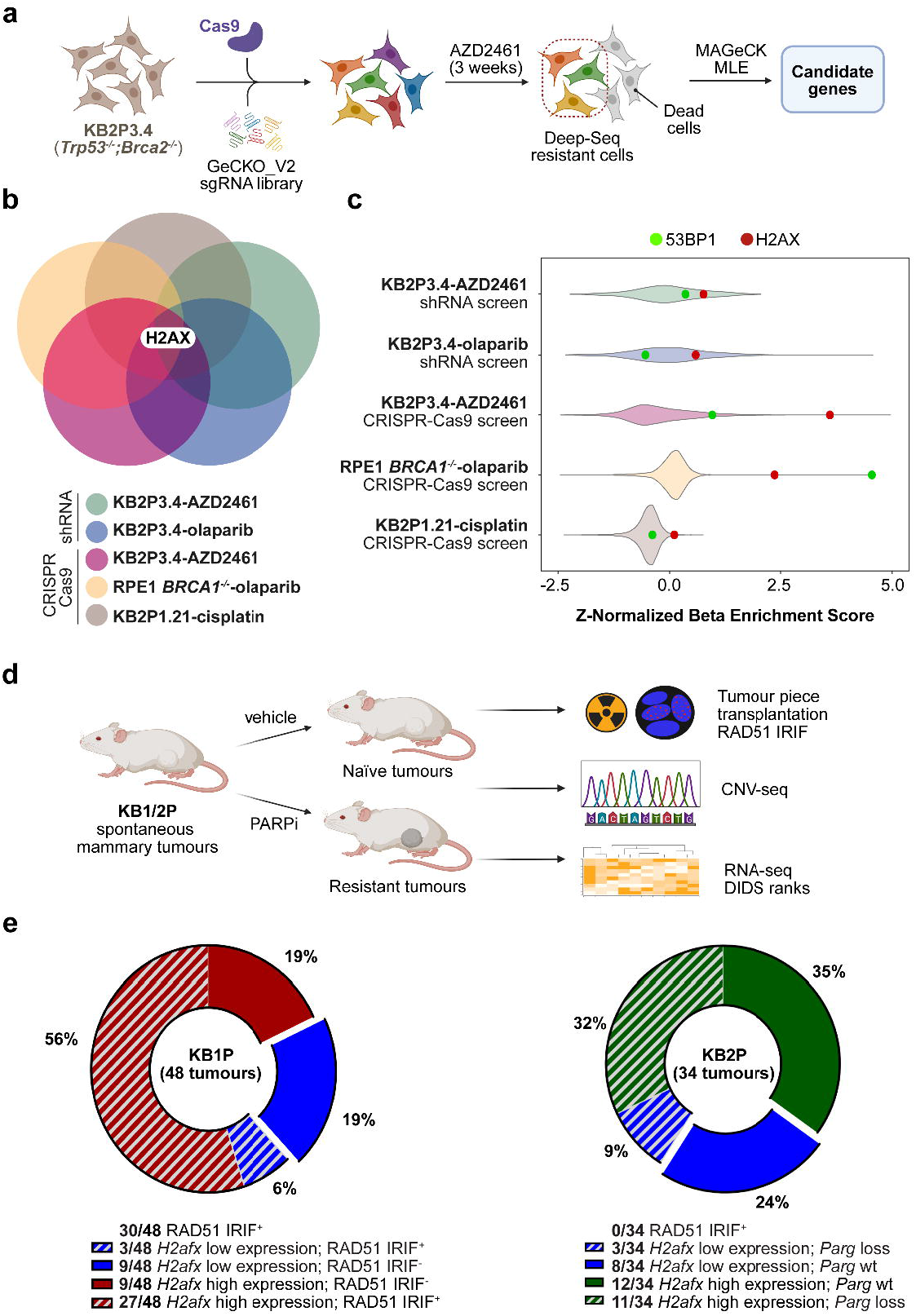
H2AX loss is frequently observed in PARPi resistant mammary tumour cells. **a**, Design of the genome-wide CRISPR-Cas9 genetic screen. **b**, Venn diagram showing the overlap of potential gene candidates identified in each individual screen. **c**, Violin plots showing the Z-Normalized Beta-Enrichment Score from 5 different genetic screens for chemoresistance carried out in different BRCA1/2-deficient cell lines. Data were all analyzed using the MaGeCK MLE algorithm to allow for cross comparison. **d**, Design of an *in vivo* pipeline to query for genetic alterations and HR restoration in PARPi-resistant mammary tumours from KB1/2P mice. **e**, Pie charts showing 48 and 34 PARPi-resistant mammary tumours from KB1P and KB2P mice, respectively. Tumours are grouped based on RAD51 IRIF^+^, *Parg* mutational status and *H2afx* gene expression (see methods text).

Considering the key role for γH2AX in DNA damage signalling^14^, it was surprising that *H2afx* loss would promote PARPi resistance. For this reason, we tested whether reduced *H2afx* gene expression would also occur in two cohorts of PARPi-resistant mammary tumours from KB1P and KB2P mice (*K14cre*;*Trp53*^*F/F*^*;Brca1*^*F/F*^ *and K14cre*;*Trp53*^*F/F*^*;Brca2*^*F/F*^, respectively), that acquired resistance *in vivo* following repeated PARPi cycles^11,15^ (Fig. 1d). In these tumours, we also determined RAD51 ionizing radiation-induced foci (IRIF) to distinguish Homologous Recombination (HR)-dependent from -independent mechanisms of PARPi resistance. We previously reported that none of the PARPi-resistant BRCA2-deficient tumours restored RAD51 IRIF^11^, whereas 63% (30/48) of the BRCA1-deficient tumours became RAD51 IRIF^+^ (Fig. 1e)^15^. Conversely, in the KB2P cohort, a large fraction of the tumours analyzed presented a major genomic structural change in the *Parg* locus, leading to the loss of *Parg* gene product (14/34) (Fig. 1e)^11^. Strikingly, we noticed that in both BRCA1/2-deficient tumour cohorts, *H2afx* gene expression was significantly reduced in a large fraction of the tumours (11/34 in the KB2P tumours and 12/48 in the KB1P tumours) with only a small overlap with *Parg* loss or HR restoration (Fig. 1e), which can be explained by the intra-tumoural heterogeneity of the resistance mechanisms^8^. Therefore, we concluded that *H2afx* downregulation frequently occurs in PARPi-resistant mammary tumours and that the underlying mechanism is likely to be different from previously reported mechanisms of resistance.

Next, we depleted *H2afx* by CRISPR-Cas9 in KB2P3.4 (*Trp53*^*-/-*^*;Brca2*^*-/-*^) and KB1P-G3 (*Trp53*^*-/-*^*;Brca1*^*-/-*^) cells and studied PARPi response *in vitro* (Extended data Fig. 1a-d). Consistent with our previous genetic analysis, *H2afx* depletion restored the cellular resistance to olaparib and AZD2461 in both KB2P3.4 and KB1P-G3 cells (Fig. 2a, b). Consistent with a toxic role for H2AX in the PARPi response in BRCA-deficient tumours, *H2afx*-depleted cells were positively selected and outgrew over the course of PARPi treatment in a competitive growth assay performed in KB2P1.21 cells (Extended Data Fig. 1e). We also generated stable *H2AFX*^*-/-*^ clones in human RPE1*-*h*TERT TP53*^*-/-*^*;BRCA1*^*-/-*^ cells and all showed full resistance to olaparib (Extended Data Fig. 1f). Importantly, we could fully complement PARPi response in KB1P-G3 cells with a wild-type H2AX but not an H2AX-S139A point mutant (Fig. 2c; Extended Data Fig. 1g). Moreover, consistent with the results from our cisplatin resistance screen (Fig. 1b, c), we also noticed that *H2afx*-deleted cells had reduced cellular sensitivity to the crosslinking agent cisplatin, but not to ionizing radiation (IR) (Extended Data Fig. 1h, i), indicating that the mechanism of resistance is likely associated with DNA replication and not general DNA damage signalling.

**Fig. 2.**
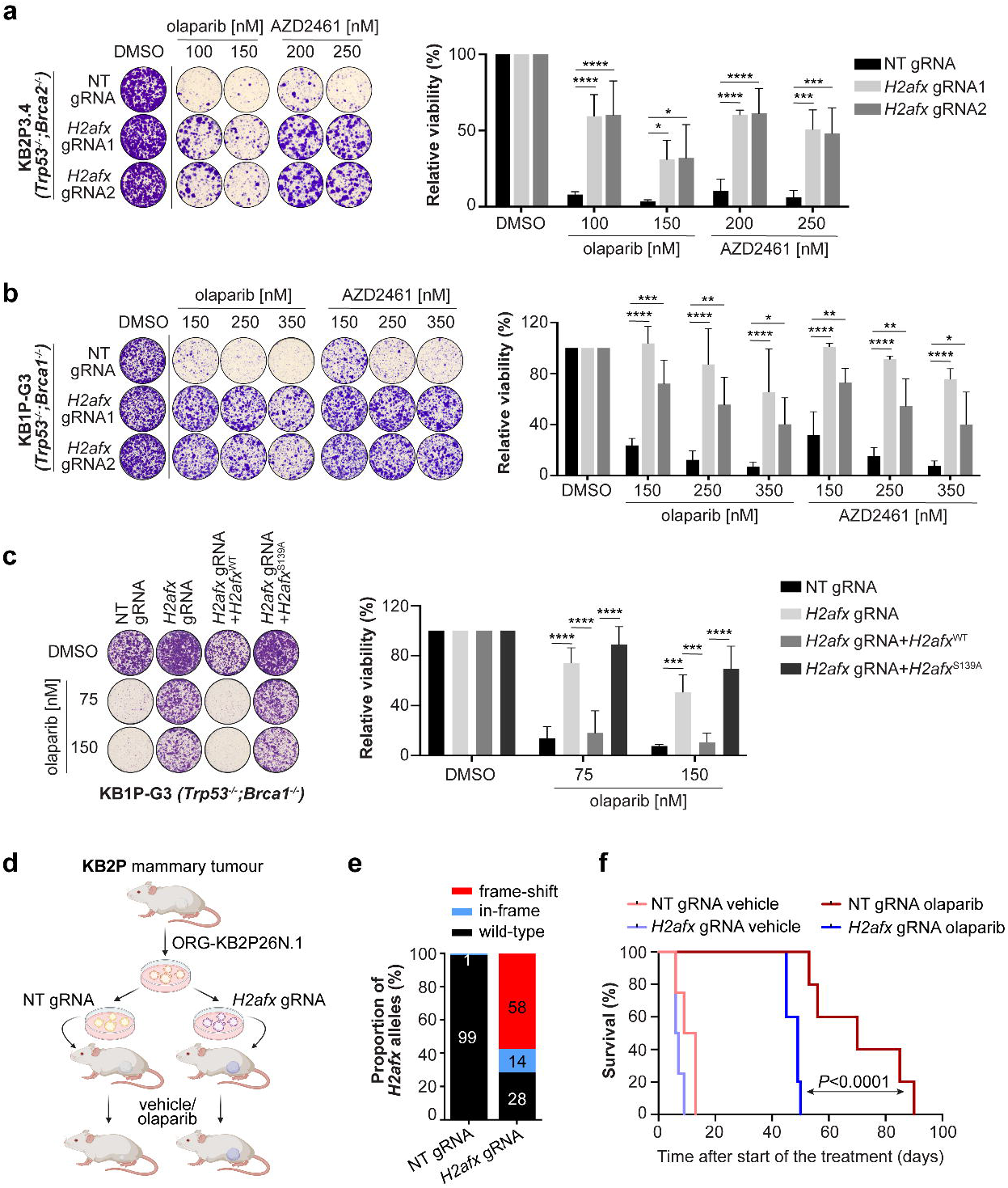
H2AX depletion leads to PARPi resistance *in vitro* and *in vivo*. **a**, Clonogenic survival assay of KB2P3.4-derived cells treated, or mock treated, with the indicated concentrations of the PARPi AZD2461 and olaparib for 12 days. Plotted values express the mean±SD of clonogenic survival (n=3). *P*-values were calculated with the unpaired two-tailed Student’s *t*-test. **b**, Clonogenic survival assay of KB1P-G3-derived cells treated as in **a. c**, Clonogenic survival assay of KB1P-G3-derived cells expressing the indicated H2AX variants and treated as in **a. d**, Schematic design of the *in vivo* experiment. **e**, Allelic modification rate of H2AX-deficient ORG-KB2P26N.1 organoids evaluated by TIDE analysis prior to transplantation. **f**, Survival of vehicle- or Olaparib-treated mice with H2AX-proficient or deficient KB2P tumours was plotted as Kaplan-Meier curve. *P*-values were calculated with the Mantel-Cox test. ^****^ *P* <0.0001.

Based on these results, we examined the impact of *H2afx* status on tumour growth *in vivo*^16^ (Fig. 2d). For this purpose, we transduced our BRCA2-deficient organoid line ORG-KB2P26N.1 with a non-targeting (NT) or a *H2afx* gRNA and verified the frameshift mutation rate by TIDE analysis (Fig. 2e). Next, we injected the modified organoids orthotopically into the inguinal mammary fat pad of mice and waited until the tumour reached a palpable size (50-100 mm^3^). Mice were then randomized and 100mg olaparib per kg or vehicle was administered by intraperitoneal injection. Strikingly, mice with *H2afx*-depleted tumours responded worse to the olaparib treatment than the *H2afx* wildtype counterparts and had to be sacrificed earlier (Fig. 2f). Together, these results confirm a crucial role for H2AX in drug resistance in BRCA-deficient tumours *in vitro* and *in vivo*.

We then dissected the molecular mechanism how H2AX loss promotes drug resistance. *H2afx*-depleted cells had reduced levels of micronuclei after olaparib treatment (Extended Data Fig. 2a), indicating reduced levels of genomic instability. Reduction in micronuclei formation is usually associated with restoration of conservative HR repair in BRCA-deficient cells^17^. Given the role of γH2AX in stabilizing 53BP1 near DNA lesions^5,6^, we tested whether H2AX loss increases PARPi resistance by preventing 53BP1 complex formation and restoring competent HR. Consistent with the function of γH2AX in recruiting 53BP1 near DNA damage sites, we observed that 53BP1 IRIF were significantly reduced in KB1P-G3 *H2afx*-depleted cells (Fig. 3a). Nevertheless, in contrast with 53BP1-deficient cells^8,13,16^, *H2afx* depletion failed to restore RAD51 IRIF (Fig. 3b), further confirming that H2AX and 53BP1 control drug resistance via two genetically separable mechanisms. We therefore hypothesized that H2AX loss increases PARPi resistance by restoring replication fork stability, another proposed mechanism of chemoresistance^18,19^. Consistent with this hypothesis, single molecule analysis of replication tracts showed that, in marked contrast with BRCA1/2-deficient cells where stalled forks undergo extensive nucleolytic degradation^20^, H2AX loss restored fork integrity in both KB1P-G3 and KB2P3.4 cells (Fig. 3c, d). Importantly, H2AX loss did not alter normal fork elongation rates nor cell growth (Extended Data Fig. 2b, c) which we instead recently observed in *Mdc1*-deleted cells^21,22^. The reduced fork resection observed in H2AX-deficient cells may be explained by a defect in the initial fork reversal^21,23–25^, or by an increased protection of the regressed arms^18,19^. To understand whether H2AX controls the fork remodeling step or the subsequent protection of the regressed arms, we directly looked at fork architecture by electron microscopy (EM) analysis^26^. Notably, we reproduced the lack of reversed fork intermediates in BRCA2-deficient cells treated with the replication fork stalling agent hydroxyurea (HU) (Fig. 3e), which has been previously attributed to the unrestrained nuclease activity at stalled forks^23–25,27^. In contrast, KB2P3.4 *H2afx*-deleted cells showed a higher number of reversed fork intermediates on the EM grids after HU treatment (Fig. 3e), indicating that H2AX is not involved in the initial reversal step but rather in the protection of reversed fork from nucleolytic degradation. Our EM analysis also revealed fewer replication intermediates with detectable single stranded DNA (ssDNA) gaps at the fork junction in KB2P3.4 *H2afx*-deleted cells (Fig. 3f), confirming a less pronounced fork degradation. MRE11 has been proposed to be the main nuclease active at stalled forks, particularly in BRCA2-deficient cells^18,23–25,27,28^ (Extended Data Fig. 2d). However, the levels of MRE11 recruitment on nascent DNA after replication stress in KB2P3.4 *H2afx*-deleted cells by *in situ* analysis of protein interactions at DNA replication forks (SIRF)^29^ were not reduced but slightly increased (Extended Data Fig. 2e). Given these results, we tested whether H2AX loss could increase replication fork protection without affecting nuclease dynamics. During the G_0_/G_1_ phase of the cell cycle, H2AX has been shown to inhibit the activity of CtIP^30^, a protein with crucial roles in DNA end resection but also with a key role in replication fork protection^31,32^. Therefore, we examined whether H2AX may similarly counteract CtIP-mediated fork protection. In agreement with previous findings^30^, we observed a stronger association of CtIP with stalled forks in KB2P3.4 *H2afx*-deleted cells (Fig. 3g). To verify whether the increased CtIP association at stalled forks reflects a functional increase in replication fork protection, we depleted CtIP function and monitored fork stability in KB2P3.4 *H2afx*-deleted cells. Since stable CtIP depletion significantly affects the proliferation of BRCA1/2-deficient cells^31^, we transiently treated KB2P3.4 *H2afx*-deleted cells with a hydrocarbon-stapled peptide (SP) that interferes with CtIP tetramerization and protein function^33^. Strikingly, pretreatment with the SP, but not with a linear peptide (LP), restored fork degradation in KB2P3.4 *H2afx*-deleted cells (Fig. 3h). We infer from these data that the effect of H2AX loss in PARPi resistance in BRCA-deficient cells is mediated by the increased association of CtIP with stalled forks.

**Fig. 3.**
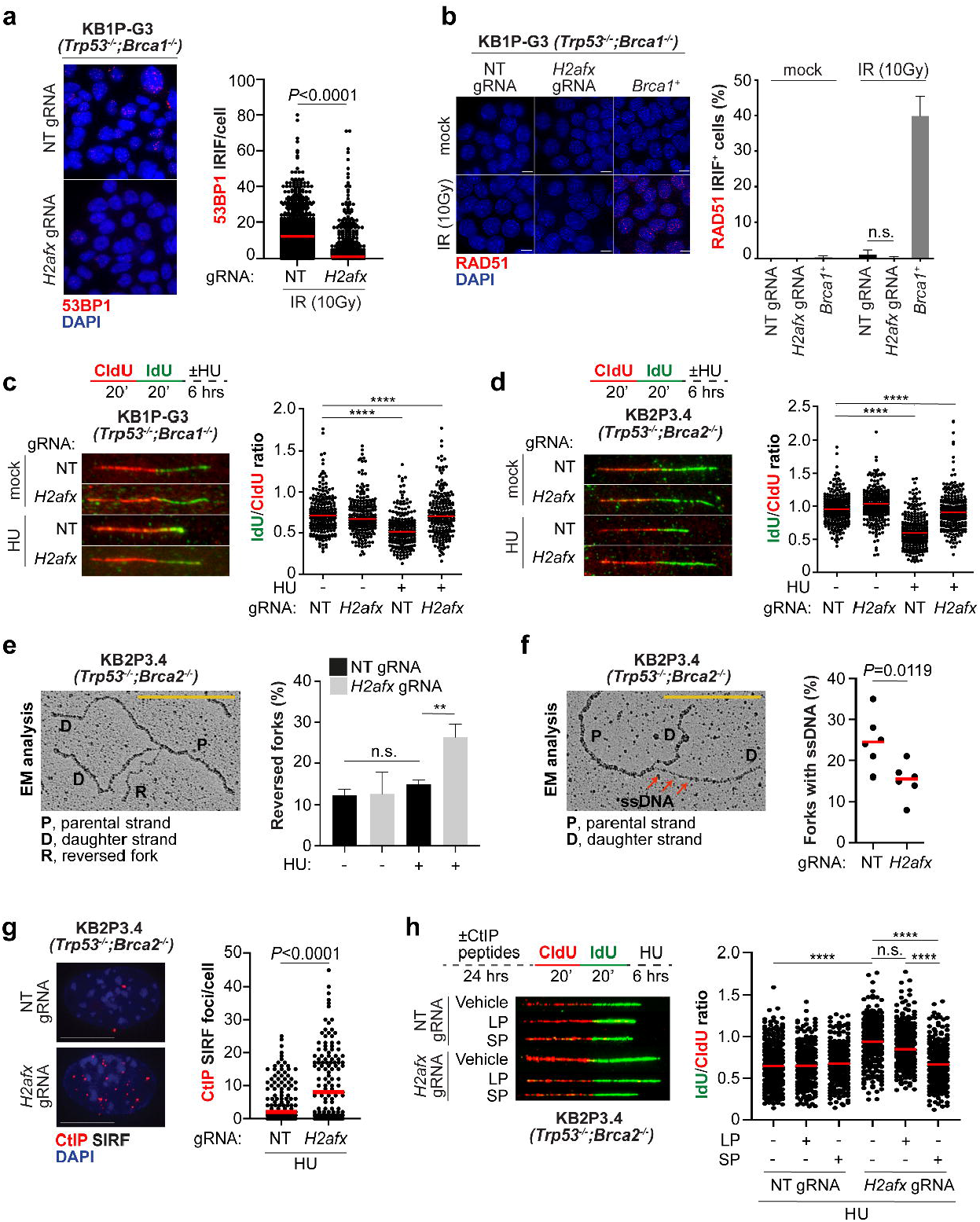
H2AX depletion restores replication fork protection. **a**, 53BP1 IRIF analysis in KB1P-G3 cells 4h after 10Gy exposure. Plotted values show the median of 53BP1 IRIF/cell from at least 600 cells (n=2). ^****^ *P*<0.0001 unpaired One-way Anova test. **b**, RAD51 IRIF analysis in KB1P-G3 cells treated as in **a**. The presented data are the mean±SD (n=3). **c**, DNA fiber analysis in KB1P-G3 cells treated according to the depicted scheme. HU was used at 8mM. Plotted values show the median of individual IdU/CldU ratios from at least 200 fibers (n=3). ^****^ *P*<0.0001 unpaired One-way Anova test. **d**, DNA fiber analysis in KB2P3.4 cells as in **c**. Plotted values show the median of individual IdU/CldU ratios from at least 200 fibers (n=3). ^****^ *P*<0.0001 unpaired One-way Anova test. **e**, EM analysis of reversed fork intermediates following HU treatment as in **c**. The electron micrograph represents a reversed replication fork. P, parental strand; D, daughter strand; R, regressed arm. The presented data are the mean±SD (n=3). *P*-values were calculated with the unpaired two-tailed Student’s *t*-test. ^**^ *P*<0.01. **f**, Dot plot shows the percentage of forks with detectable ssDNA gaps, as exemplified in the depicted micrograph. *P*-values were calculated with the unpaired two-tailed Student’s *t*-test. ^**^ *P*<0.01. **g**, CtIP SIRF in KB2P3.4-derived cells treated as in **c**. Plotted values show the median of CtIP SIRF foci/cell from at least 150 cells (n=3). *P*-values were calculated with the unpaired two-tailed Student’s *t*-test. ^****^ *P*<0.0001. **h**, KB2P3.4-derived cells were pretreated for 24h with the indicated CtIP peptides prior to analog *in vivo* labeling according to the depicted scheme. HU was used at 8mM and the CtIP peptides were used at 10μM. Plotted values show the median of individual IdU/CldU ratios from at least 250 fibers (n=2). ^****^ *P*<0.0001 unpaired One-way Anova test.

Chemoresistant tumours often develop new genetic vulnerabilities that can be exploited in targeted therapies^34^. While searching for new genetic vulnerabilities in H2AX-deficient tumours, we have found that *H2AFX* scored as one of the strongest hits in loss-of-function CRISPR-Cas9 screens performed in distinct cancer cell lines treated with the ATMi AZD0156^35^ (Fig. 4a, b). Therefore, we hypothesized that ATMi may restore PARPi sensitivity in our KB1P-G3 and KB2P3.4 *H2afx*-deleted cells. Consistent with this idea, different AZD0156 doses (5-100nM) efficiently restored olaparib sensitivity in both KB1P-G3 and KB2P3.4 *H2afx*-deleted cells (Fig. 4c; Extended Data Fig. 3a). ATM is known to phosphorylate and activate CtIP at DSBs during DNA end resection^36^ (Extended Data Fig. 3b). We therefore investigated whether ATM inhibition restores PARPi sensitivity by interfering with CtIP-mediated fork protection. Indeed, pretreatment with the ATMi AZD0156 robustly restored fork degradation in KB2P3.4 *H2afx*-deleted cells (Fig. 4d). To confirm that this effect was due to the lack of CtIP regulation and not due to the lack of phosphorylation of other ATM targets, we generated doxycycline-inducible U-2OS lines that express either a siRNA-resistant CtIP^WT^ or a CtIP^8A^ variant where all the S/T-Q sites were mutated to alanine (Extended Data Fig. 3c). Strikingly, CtIP^WT^ expression complemented the lack of fork protection after endogenous CtIP depletion by siRNA, while the CtIP^8A^ variant failed to do so (Fig. 4e). This shows that ATM-mediated CtIP phosphorylation is critical for fork protection.

**Fig. 4.**
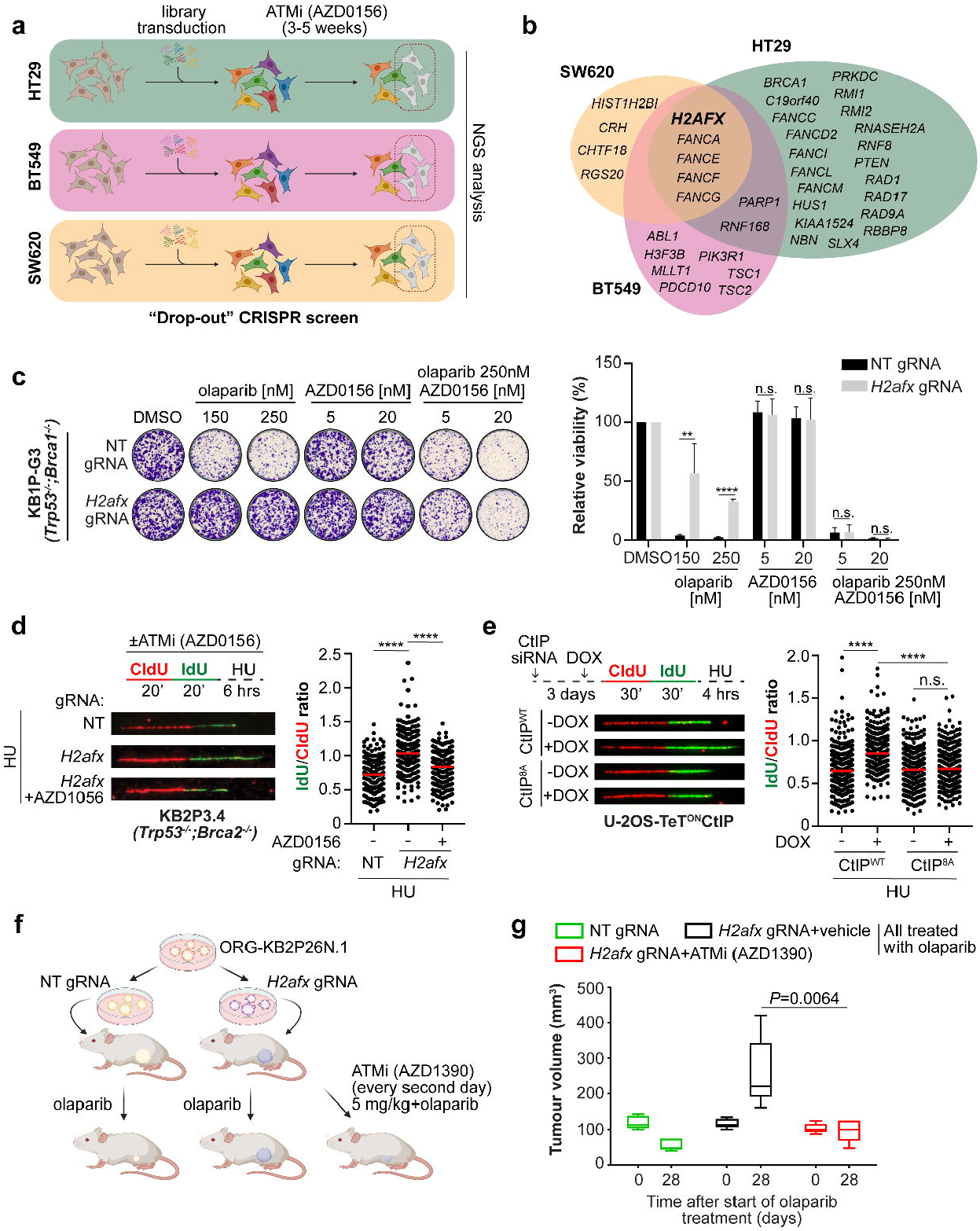
ATM inhibition restores PARPi sensitivity in H2AX-deficient mammary tumours. **a**, Design of the functional CRISPR-Cas9 drop-out screen with the ATMi AZD0156 in distinct cancer cell lines. **b**, Venn diagram of the top hits from each individual CRISPR-Cas9 screen. **c**, Clonogenic survival assay of KB1P-G3-derived cells treated or mock with the indicated concentrations of olaparib with and without AZD0156 for 12 days. Plotted values express the mean±SD of clonogenic survival (n=3). *P*-values were calculated with the unpaired two-tailed Student’s *t*-test. ^**^*p*<0.01, ^****^*p*<0.0001. **d**, DNA fiber analysis in KB2P3.4-derived cells treated according to the depicted scheme. HU was used at 8mM and AZD0156 was used at 10μM. Plotted values show the median of individual IdU/CldU ratios from at least 170 fibers (n=3). ^****^ p<0.0001 unpaired One-way Anova test. **e**, DNA fiber analysis in U-2OS-derived cells transfected with CtIP siRNA and treated according to the depicted scheme. 24h before the experiment, CtIP^WT^ or CtIP^8A^ expression was induced by doxycycline. Plotted values show the median of individual IdU/CldU ratios from at least 300 fibers (n=2). ^****^ p<0.0001 unpaired One-way Anova test. **f**, Schematic design of the *in vivo* experiment with the ATMi AZD1390. **g**, Plotted values express the size (mm^3^) of individual tumours from animals transplanted with the indicated organoid lines and subjected to the indicated treatment for 28 consecutive days after tumor formation. P value was calculated with the two-way Anova.

Moreover, we explored whether ATMi restores PARPi sensitivity also in H2AX-deficient tumours *in vivo* (Fig. 4f). To address this question, we performed a similar experiment as shown in Fig. 2e and added a group of mice treated with the new and clinically relevant ATMi AZD1390^37^ (5 mg/kg, every second day) in combination with olaparib for the whole duration of the experiment (Fig. 4f). Importantly, at this dosage of AZD1390, the animals did not show any undesirable effects such as reduction of body weight (data not shown). Again, we observed that in the animals with the *H2afx*-depleted organoids, tumours did not respond to the olaparib treatment and grew over the therapy course compared to the tumours in the control group (Fig. 4g). However, consistent with our *in vitro* data, the AZD1390 combination restored PARPi response and impaired tumour growth throughout the duration of the experiment (Fig. 4g). Together, these data indicate that ATM inhibitors in combination with PARPi can be effectively used to overcome drug resistance in H2AX-deficient tumours and control tumor growth.

In conclusion, we have shown a new role for the histone H2AX in replication fork biology in BRCA1/2-deficient tumours. Previous work in BRCA-proficient cells proposed a function for the histone H2AX in maintaining the stability of reversed replication forks during unstressed DNA replication^38^. Interestingly, the latter function appeared to depend on ATM-mediated phosphorylation of Serine 139 and the RNF8-RNF168 axis^38^. Here, with our molecular characterization of replication fork dynamics and fork architecture in BRCA1/2-deficient H2AX-deficient cells, we have now found that H2AX loss restores replication fork stability in BRCA-deficient tumours and increases chemotherapy resistance without restoring homology-directed DNA repair capacity. Our discovery may have implications for the treatment of *BRCA*-mutated cancers and contribute to our understanding of mechanisms of resistance that are independent of restoration of BRCA1/2 function^39^. These findings also shed new lights on the mechanisms leading to cross-resistance between PARPi and platinum-derived therapies, which often occurs in *BRCA*-mutated high-grade serous ovarian carcinoma patients. Moreover, it shows that the newly identified role of H2AX in replication fork degradation is crucial for the efficacy of DNA replication-targeting drugs in *BRCA*-mutated tumours.

## Methods

### Cell culture conditions

KB2P3.4 (*Trp53*^*-/-*^*;Brca2*^*-/-*^), KB2P1.21 (*Trp53*^*-/-*^*;Brca2*^*-/-*^), and KB1P-G3 (*Trp53*^*-/-*^*;Brca1*^*-/-*^) cells were derived from KB2P and KB1P mammary tumours as previously described^7,8^. KB1P1.21 *Mdc1*^*-/-*^ cells were generated as recently reported^21,22^. All mouse-derived cells were grown at 37ºC in 3% O_2_ and cultured in Dulbecco’s modified Eagle medium Nutrient mixture F-12 (DMEM/F12; Gibco) supplemented with 10% Fetal Bovine Serum (FBS), pen/strep solution (50 U/ml), 5 ng/ml cholera toxin (Sigma Aldrich), insulin (5 μg/ml, Sigma Aldrich) and 5 ng/ml murine Epidermal Growth Factor (mEGF, Sigma Aldrich). ORG-KB2P17S.1 organoids were grown at 37ºC in normal oxygen embedded in Cultrex Reduced Growth Factor Basement Membrane Extract Type 2 (BME, Trevigen), seeded on 24-well suspension plates (Greiner Bio-One) and cultured in complete mouse mammary gland organoid medium: AsDMEM/F12 supplemented with 1M HEPES (Sigma), GlutaMAX (Invitrogen), pen/strep (Gibco), B27 (Gibco), 125μM N-acetyl-L-cysteine (Sigma), 50 ng/ml murine epidermal growth factor (mEGF, Invitrogen). Human RPE1-h*TERT TP53*^*-/-*^, RPE1-h*TERT TP53*^*-/-*^*;BRCA1*^*-/-*^, and the RPE1-h*TERT TP53*^*-/-*^*;BRCA1*^*-/-*^;*H2AFX*^*-/-*^ clones were grown at 3% O_2_ in Dulbecco’s modified Eagle medium Nutrient mixture F-12 (DMEM/F12; Gibco) and penicillin/streptomycin solution (100 U/ml). SW62O cells were grown in L-15 medium (Leibovitz), 10% FBS, 1X glutamax. HT29 cells were grown in EMEM, 10% FBS, 1% non-essential aminoacids, 1% L-glutamine. BT549 cells were grown in RPMI-1640, 10% FBS, 0.023 U/ml insulin. U-2OS-TeT^ON^-eGFP-CtIP^WT^ and U-2OS-TeT^ON^-eGFP-CtIP^8A^ were grown at 37ºC in DMEM supplemented with 10% FCS, 1% pen/strep, 5 μg/ml Blasticidin, 200 μg/ml Zeocin. CtIP expression was induced 24h before the experiment with 1 μg/ml of doxycycline.

All cell lines were authenticated by *Brca1/2*-specific PCR-based genotyping (mouse)^7,8^ and they were regularly tested for mycoplasma contamination (Mycoalert, Lonza).

### Drugs and reagents

The following chemical reagents were used throughout the study: AZD2461 (kindly provided by AstraZeneca), olaparib (kindly provided by AstraZeneca and Syncom (Groningen, The Netherlands)), cisplatin (Teva; #7680479980428), Hydroxyurea (Sigma; #H8627), Mitomycin C (Sigma; #M4287), AZD1056 (kindly provided by AstraZeneca), AZD1390 (kindly provided by AstraZeneca), Mirin (Sigma; #M9948), 5*-*Iodo-2’-deoxyuridine (IdU) (Sigma; #I7125), 5-Chloro-2’-deoxyuridine (CldU) (Sigma; #C6891). LP and SP (Bachem; #4143690 and #4111111, respectively).

### CRISPR/Cas9-based genetic screens

The PARPi resistance screens were performed in the KB2P3.4 tumor cell line, which was previously established from a KB2P tumor^7^. Mouse GeCKO_V2 library, pool B (62,804 gRNAs targeting 20,628 genes (3 gRNAs/gene) including 1,000 non-targeting gRNAs), was stably introduced into the cells by lentiviral transduction at multiplicity of infection (MOI) of 1.5. Mouse GeCKO_V2 CRISPR knockout pooled library was a kind gift from Feng Zhang^9^. 6 independent transductions were carried out to obtain mutagenized cells for biological replicates of the PARPi resistance screen. To perform the genetic screen with a 100x library coverage, 6×10^6^ mutagenized KB2P3.4 cells in each replicate were plated in 10-cm flasks, at low density (30,000 cells per flask) and grown in medium containing 200nM AZD2461 for 3 weeks. The medium with the PARPi was refreshed twice a week. Cells were harvested before and after PARPi treatment for genomic DNA isolation. Subsequently, gRNA sequences were amplified from genomic DNA by two rounds of PCR amplification^40^. Resulting PCR products were purified using MinElute PCR Purification Kit (Qiagen) and submitted for Illumina sequencing. Quality control was performed using R software (R Core Team, 2022) and package *edgeR*^41^. Sequence alignment and enrichment analysis (day 0 vs PARPi-treated population) was carried out using the R package *MAGeCKFlute*^42^. Dataset of MAGeCK MLE analysis results of the CRISPR/Cas9 screen on RPE1-h*TERT* cells was extracted from the Supplementary Table 1 of Nordermeer et al., 2018^12^.

For the CRISPR-Cas9 screens with the ATMi AZD1056, SW620, HT29, and BT549 cells were infected with lentiviral particles containing the whole-genome sgRNA library (Horizon Discovery), subjected to puromycin selection, and passaged to ensure loss of affected protein products. Puromycin-resistant cells were exposed to 10□nM ATMi (AZD0156) for 21 days (SW620, HT29) or 35 days (BT549), and remaining cell pools were isolated. Genomic DNA was extracted from these (QIAamp DNA Blood Maxi kit (Qiagen #51194)) and from parallel cell cultures treated in the absence of AZD0156, and DNA libraries were prepared and sequenced using an Illumina NextSeq next generation sequencing platform. Analysis of NGS data sets *e*.*g*. sgRNA abundance was achieved using Horizon Discovery’s data processing scripts, based on published analysis tool MAGeCK.

### DDR shRNA-based genetic screens

PARPi resistance shRNA screens were previously described^11^. Briefly, a shRNA library targeting a DNA Damage Response (DDR) gene set was built based on a gene list described before and the NCBI search (terms: “DNA repair”, “DNA damage response”, “DNA replication”, “telomere-associated genes”)^40^. The shRNA library was stably introduced into the tumor cell line KB2P3.4, which was subsequently selected with the PARPi AZD2461 or olaparib for 3 weeks. Genomic DNA was purified before and after treatment, amplified and sequenced as described above. Sequence alignment and analysis were performed using the MAGeCK software^10^, MAGeCK-VISPR Maximum Likelihood Estimation (MLE) module, and the R package *MAGeCKFlute*^42^.

### Lentiviral transductions

Lentiviral stocks were generated by transient transfection of HEK293FT cells. On day 0, 6×10^6^ HEK293FT cells were seeded in 150-cm cell culture dishes and on the next day transiently transfected with lentiviral packaging plasmids and the pLentiCRISPRv2 vector containing the respective *H2afx*-targeting gRNA or a non-targeting gRNA using 2x HBS (280nM NaCl, 100mM HEPES, 1.5mM Na_2_HPO_4_, pH 7.22), 2.5M CaCl_2_ and 0.1x TE buffer (10mM Tris pH 8.0, 1 mM EDTA pH 8.0, diluted 1:10 with dH_2_O). After 30h, virus-containing supernatant was concentrated by ultracentrifugation at 20,000 rpm for 2h in a SW40 rotor and the virus pellet was finally resuspended in 100μl PBS. The virus titer was determined using a qPCR Lentivirus Titration Kit (#LV900, Applied Biological Materials). For lentiviral transduction, 150,000 target cells were seeded in 6-well plates. 24h later, virus at the MOI of 50 was applied with 8 μg/ml Polybrene (Merck Millipore). Virus-containing medium was replaced with medium containing puromycin (3.5 μg/ml, Gibco) 24h later. Puromycin selection was performed for three days; subsequently cells were expanded and frozen down at early passage. Tumor-derived organoids were transduced according to a previously established protocol^16^. The target sites modifications of the polyclonal cell pools were analyzed by TIDE analysis.

### Genome editing

For CRISPR/Cas9-mediated genome editing, KB2P3.4 and KB2P1.21 cells or KB2P17S.1 tumor-derived organoids were transduced with the pLentiCRISPRv2 vector encoding non-targeting gRNA, *H2afx*-targeting gRNA1 or *H2afx*-targeting gRNA2. The cells were then grown under Puromycin (3 μg/ml) selection for 5 days. All constructs were verified with Sanger sequencing. For CRISPR/Cas9-mediated targeting of *H2afx* gene in KB1P-G3 cells, non-targeting gRNA or *H2afx*-targeting gRNA1 or gRNA2 were cloned into the pX330 vector (Addgene; #42230). Sanger sequencing-verified pX330 plasmids containing the correct sequences of gRNAs were transfected into KB1P-G3 cells using the TransIT-LT1 transfection reagent (Mirus) according to the manufacturer’s protocol. The cells were then grown under Puromycin (3 μg/ml) selection for 5 days. CRISPR gRNA sequences for *H2afx* modification were chosen from the GeCKO_V2 library. The gRNA sequences were as follows: m*H2afx* gRNA1: 5’- TCGTACACTATGTCCGGACG -3’; m*H2afx* gRNA2: 5’- GGCGCCGGCGGTCGGCAAGA -3’; Non-Targeting (NT) gRNA: 5’- TGATTGGGGGTCGTTCGCCA-3’; h*H2AFX* gRNA1: 5’-GACAACAAGAAGACGCGAATC- 3’. H2ax reconstitution was performed using the pOZ-N-FH (a kind gift from Dipanjan Chowdury). The *H2afx* coding sequence from *Mus musculus* was ordered from Eurofins and cloned into the pOZ-N-FH backbone adding the 1xHA tag at the N terminus using the in-fusion HD cloning kit (#12141, Takara). Full length wild type *H2afx* coding sequence was then mutagenized to obtain the desired S139A point mutation.

### gDNA isolation, amplification, and TIDE analysis

To assess the modification rate at the gRNA-targeted region of *H2afx*, cells were pelleted, and genomic DNA was extracted using the QIAmp DNA mini kit (Qiagen) according to manufacturer’s protocol. Target loci were amplified using Phusion High Fidelity Polymerase (ThermoFisher Scientific) using a 3-step protocol: 98°C for 30’’, 35 cycles at 95°C for 15’’, 55°C for 15’’and 72°C for 30’’, 72°C for 7’. Reaction mix consisted of 10μl of 2x Phusion Mastermix (ThermoFisher Scientific), 1μl of 10μM forward (5’-CAATCACTGGGCGCGTTC-3’) and reverse (5’- TGGCTCAGCTCTTTCTGTGAG-3’) primers and 100 ng of DNA in 20μl total volume. PCR products were purified using the QIAquick PCR purification kit (Qiagen) according to manufacturer’s protocol and submitted with corresponding sequencing primers for Sanger sequencing to confirm target modifications using the TIDE algorithm. The sequencing primers used for analysis of the modification rate in mouse cells are the following: mgRNA1 seq. primer: 5’- CAATCACTGGGCGCGTTC-3’; gRNA2 seq. primer: 5’-GAGTACCTCACTGCCGAG-3’; hgRNA1 seq. primer: 5’-GACAACAAGAAGACGCGAATC-3’.

### *H2afx* gene expression analysis in PARPi-resistant mammary tumours

Differential *H2afx* gene expression analysis from matched PARPi naïve and resistant mammary tumours was performed as previously described using a threshold of *p*<0.05 for statistical significance^11^.

### Clonogenic assays

To assess the growth and survival upon treatment with PARPi, cisplatin or irradiation, KB2P3.4 and KB1P-G3 cells were seeded in 6-well plates in the following densities: 3,000 cells/well (KB2P3.4) and 4,000 cells/well (KB1P-G3). The treatment of cells with DMSO or indicated concentrations of PARPi olaparib or AZD2461 started at the day of plating the cells and lasted for the whole duration of the experiment. The medium with DMSO or PARPi was refreshed twice a week. The control, DMSO-treated plates were fixed 7 days after seeding, the PARPi-treated plates were fixed after 10 days. For the cisplatin treatment, cells were plated 24h prior addition of cisplatin-containing media. After 24h, medium was refreshed and all the plates were fixed after 7 days. Irradiation was carried out in a fractionated manner using the indicated irradiation doses 24, 48 and 72h following plating of the cells. Plates with non-irradiated cells were fixed 7 days post plating, the irradiated cells were fixed after 10 days. The fixation was done with 4% formalin and the surviving colonies stained with 0.1% crystal violet. The cell survival and growth were analyzed in an automated manner using the ImageJ ColonyArea plugin. For the competition assays, cells were collected before and after the experiment for gDNA isolation and TIDE analysis as described above. For clonogenic assays with ATMi, the same experimental setup as in the clonogenic assays with PARPi was used. ATMi was added to the medium on the day of plating and the medium containing the drug was refreshed twice/week until the end of the experiment.

### *In vivo* study

All animal experiments were approved by the Animal Ethics Committee of The Netherlands Cancer Institute (Amsterdam, the Netherlands) and the Canton of Bern (license BE69/2021) and were performed in full compliance with national laws, which enforce Dir. 2010/63/EU (Directive 2010/63/EU of the European Parliament and of the Council of 22 September 2010 on the protection of animals used for scientific purposes). For tumor organoid transplantation, ORG-KB2P26N.1 organoids were collected, incubated with TripLE at 37°C for 5’, dissociated into single cells, washed in PBS, resuspended in tumor organoid medium, and mixed in a 1:1 ratio of tumor organoid suspension and BME. Organoid suspension containing a total of 10^5^ cells were injected in the fourth right mammary fat pad of 6–9-week-old NMRI nude mice. Mammary tumor size was measured by caliper and tumor volume was calculated ((length x width^2^) /2). Treatment of tumor bearing mice was initiated when tumours reached a detectable size of at least ∼25-50 mm^3^, at which point mice were separated into a vehicle-treated group (NT gRNA n= 5, *H2afx*-targeting gRNA1 n= 5) and an olaparib-treated group (NT gRNA n= 5, *H2afx*-targeting gRNA n= 5). Olaparib (100 mg/kg) was administered by intraperitoneal injections for 28 consecutive days. The control tumor-bearing mice were dosed with vehicle following the same the schedule as the PARPi group. Animals were anesthetized with isoflurane, sacrificed with CO_2_ followed by cervical dislocation when the tumor reached a volume of approximately 1,000 mm^3^. Tumor sampling included cryopreserved tumor pieces, fresh frozen tissue, and formalin-fixed material (4% (v/v) formaldehyde in PBS).

### Immunofluorescence

Cells were seeded on coverslips in 24-well plates 3 days prior the experiment. To analyze 53BP1 and RAD51 foci formation in H2AX-deficient KB1P-G3 cells, DNA damage was induced by γ-irradiation (10Gy) 4h prior to fixation. Subsequently, cells were washed in PBS and fixed with 4% (v/v) PFA/PBS for 20’ at RT. Fixed cells were washed with PBS and permeabilized for 20’ in 0.2% (v/v) Triton X-100/PBS. Next, slides were washed three times with 0.2% Tween-20/PBS and blocked with staining buffer (PBS, BSA (2% w/v), glycine (0.15% w/v), Triton X-100 (0.1% v/v)) for 1h at RT. Incubation with the primary rabbit polyclonal anti-53BP1 (#A300-272A, Bethyl laboratories) and anti-RAD51 antibody (#70-012, Bioacademia) diluted 1:1000 in staining buffer was carried out for 2h at RT. Slides were then washed four times for 5’ with 0.2% (v/v) PBS-Tween-20 and then incubated with Goat anti-rabbit IgG (H+L) Cross-Absorbed Secondary Antibody, Texas Red-X (# T-6391, ThermoFisher Scientific) diluted 1:2000 in staining buffer for 1h at RT. Slides were washed three times for 5’ with 0.2% PBS-Tween-20, once with PBS and then mounted with Duolink *In Situ* mounting medium with DAPI (#DUO82040, Sigma Aldrich). Z-stack fluorescent images were acquired using the DeltaVision Elite widefield microscope (GE Healthcare Life Sciences). Multiple fields of view were imaged per sample with Olympus 100X/1.40, UPLS Apo, UIS2, 1-U2B836 objective and sCMOS camera at the resolution 2048 × 2048 pixels. Deconvolution of the acquired images was performed by the softWoRx DeltaVision software. Image analysis was performed using Fiji image processing package of ImageJ. Briefly, all nuclei were detected by the “analyze particles” command and all the foci per nucleus were counted with the “find maxima” command. Data were plotted with Prism software.

### Analysis of micronuclei formation

KB2P3.4 cells were seeded on coverslips in 24-well plates and treated with DMSO or indicated concentrations of olaparib 24h later. After 48h of treatment, cells were washed with PBS and fixed with 4% (v/v) PFA/PBS for 20’ in RT. Cells were then washed 3 times in 0.2% (v/v) PBS-Tween-20 and permeabilized for 20’ in 0.2% (v/v) Triton X-100/PBS. Subsequently, slides were washed 3 times with PBS, counterstained with DAPI (1:50,000 dilution, #D1306, Life Technologies) and washed 5 more times with PBS before mounting in Fluorescence mounting medium (#S3023, Dako). Z-stack images were acquired using the DeltaVision Elite widefield microscope (GE Healthcare Life Sciences). Multiple fields of view were imaged per sample with Olympus 100X/1.40, UPLS Apo, UIS2, 1-U2B836 objective and sCMOS camera. The frequency of micronuclei positive cells was analyzed manually in Fiji.

### Replication fork progression by DNA fiber analysis

Fork progression was measured as described previously^43^. Briefly, asynchronously growing subconfluent KB2P1.21 or KB2P3.4 cells were labeled with 30μM thymidine analogue 5-chloro-2’-deoxyuridine (CIdU) (#C6891, Sigma-Aldrich) for 20’, washed three times with warm PBS and exposed to 250μM of 5-iodo-2′-deoxyuridine (IdU) for 20’. All cells were collected by trypsinization and 2μl of this cell suspension was then mixed with 8μL of lysis buffer (200mM Tris-HCl, pH 7.4, 50mM EDTA, and 0.5% (v/v) SDS) on a positively charged microscope slide. After 9’ of incubation at RT, the slides were tilted at an approximately 30-45° angle to stretch the DNA fibers onto the slide. The resulting DNA spreads were air-dried, fixed in 3:1 methanol/acetic acid, and stored at 4°C overnight. Next day, the DNA fibers were denatured by incubation in 2.5M HCl for 1h at RT, washed five times with PBS and blocked with 2% (w/v) BSA in 0.1% (v/v) PBST (PBS and Tween 20) for 40’ at RT while gently shaking. The newly replicated CldU and IdU tracks were stained for 3h at RT using two different anti-BrdU antibodies recognizing CldU (#ab6326, Abcam) and IdU (#347580, BD Biosciences), respectively. After washing five times with PBS-T the slides were stained with goat the anti-mouse IgG (H+L) Cross-Adsorbed Secondary Antibody, Alexa Fluor 488 (#A-11029, ThermoFisher Scientific) diluted 1:600 in blocking buffer and with the Cy3 AffiniPure F(ab’)□ Fragment Donkey Anti-Rat IgG (H+L) antibody (#712-165-513, Jackson Immuno Research) diluted 1:150 in blocking buffer. Incubation with secondary antibodies was carried out for 1h at RT in the dark. The slides were washed five times for 3’ in PBS-T, air-dried and mounted in Fluorescence mounting medium (#S3023, Dako). Fluorescent images were acquired using the DeltaVision Elite widefield microscope (GE Healthcare Life Sciences). Multiple fields of view from at least two slides (technical replicates) of each sample were imaged using the Olympus 60X/1.42, Plan Apo N, UIS2, 1-U2B933 objective and sCMOS camera at the resolution 2048 × 2048 pixels. To assess fork progression, the sum of individual CldU and IdU track lengths was measured using the segmented line tool in ImageJ software. Statistical analysis was carried out using Prism.

### Replication fork stability by DNA fiber analysis

CldU and IdU pulse-labeling of asynchronously growing KB2P3.4 and KB1P-G3 cells expressing NT or *H2afx*-targeting gRNA was performed as described above. After pulse-labeling with IdU and three washes with PBS, medium containing 8mM hydroxyurea (HU) was added for 6h. Cells were then washed and harvested by trypsinization, and then processed as described above. Replication fork stability was analyzed by measuring the ratio between CldU and IdU tracks in ImageJ.

### Immunoblotting

Cells were lysed for 40’ in RIPA buffer supplemented with Halt Protease and Phosphatase Inhibitor Cocktail (100x) (#78420, Thermo Fisher Scientific) while briefly vortexed every 10’. Lysates were then centrifuged at 10,000 rpm for 10’ at 4°C and the supernatant was collected to determine protein concentration using Pierce BCA Protein Assay Kit (#23225, Thermo Fisher Scientific). Before loading, protein lysates were denatured at 95°C for 5’ in 6x SDS sample buffer. Proteins were separated by SDS/PAGE in 10% gel before wet transfer to 0.45μm nitrocellulose membranes (GE Healthcare) and blocked in 5% dry milk powder in TBS-T (100mM Tris, pH 7.5, 0.9% NaCl, 0.05% Tween-20). Membranes were incubated with the mouse monoclonal anti-HA (1:1,000, #901533, Biolegend) and anti-γ-Tubulin (1:1,000, #5886, Cell Signalling), anti-GFP (1:1,000, #2555, Cell Signalling) primary antibodies for 2h at RT. After three 5’ washes in TBS-T, anti-rabbit or anti-mouse Horseradish Peroxidase (HRP)-linked secondary antibodies (1:5,000, Cell Signalling) were applied for 1h at RT. Images were acquired using Vilber FUSION FX chemiluminescent imager.

### SIRF (*in Situ* analysis of protein Interactions at DNA Replication Forks)

SIRF assay was performed as previously reported^29^. Cells were seeded on coverslips and the following day they were pulsed with 25μM EdU for 10’. After the EdU pulse, cells were initially pre-extracted with CSK buffer on ice for 5’ and then fixed with 3.7% Paraformaldehyde at RT for 10’. Coverslips were then washed with PBS and stored overnight at 4ºC. The following day cells were permeabilized in 0.2% Triton X-100 in PBS for 5’ and then the click reaction (100mM Tris pH 8, 100mM CuSO_4_, 2mg/ml sodium-L-ascorbate, 10mM biotin-azide) was performed for 90’ at 37ºC. Slides were then blocked for 1h at 37ºC with blocking solution (PBS, BSA 2%, glycine 0.15%, Triton X-100 0.1%), followed by incubation with primary antibodies for 1h at 37ºC (rabbit anti-CtIP 1:500, Bethyl Laboratories #A300-487A; mouse anti-Biotin 1:200, #200-002-211, Jackson Immuno Research; rabbit anti-MRE11 1:500, a kind gift from Arnab Ray Chaudhuri). After antibody incubation, coverslips were washed 2X with Buffer A for 5’ at RT (Duolink kit). Each coverslip was then incubated for 1h at 37ºC with Duolink PLA probes (Thermo Fisher Scientific) diluted in blocking solution. After 2 washes with Buffer A for 5’ at RT, probes were ligated for 30’ at 37ºC and amplified by polymerase reaction for 100’ at 37ºC. Coverslips were then washed 2X with Buffer B for 5’ at RT (Duolink kit) and then mounted with DAPI on microscope slides. Images were acquired on multiple stacks using the DeltaVision Elite widefield microscope with a 60X objective. Deconvolution of the images was done using the softWoRx DeltaVision software. The number of foci in each cell was scored with ImageJ and the statistical analysis was performed using Prism.

### Transmission electron microscopy of replication intermediates

The procedure was performed as described previously with minor modifications^26^. A total of 2.5–5.0×10^6^ asynchronously growing KB2P3.4 cells expressing either NT or *H2afx*-targeting gRNA were treated with 8mM hydroxyurea for 5h, washed with PBS and then harvested by trypsinization and resuspended in 10mL of cold PBS. DNA was cross-linked by exposing the living cells twice to 4,5′,8-trimethylpsoralen at a final concentration of 10 μg/mL followed by 3’ irradiation pulses with UV 365-nm monochromatic light (UV Stratalinker 1800, Agilent Technologies). The cells were then washed repeatedly with cold PBS and lysed with a cell lysis buffer (1.28M sucrose, 40mM Tris-Cl, pH 7.5, 20mM MgCl_2_, and 4% (v/v) Triton X-100). The nuclei were then digested in a digestion buffer (800mM guanidine-HCl, 30mM Tris-HCl, pH 8.0, 30mM EDTA, pH 8.0, 5% (v/v) Tween 20, and 0.5% (v/v) Triton X-100) supplemented with 1 mg/mL proteinase K at 50°C for 2h. Genomic DNA was extracted with a 24:1 Chloroform:Isoamyl alcohol mixture by phase separation (centrifugation at 8,000 rpm for 20’ at 4°C) and precipitated by addition of equal amount of isopropanol to the aqueous phase, followed by another centrifugation step (8,000 rpm for 10’ at 4°C). The obtained DNA pellet was washed once with 1mL of 70% ethanol, air-dried at RT, and resuspended by overnight incubation in 200μL TE (Tris-EDTA) buffer at RT. 12μg of the extracted genomic DNA was digested for 5h at 37°C with 100U restriction enzyme *PvuI*I-HF (#R3151S, New England Biolabs). The digest was cleaned up using a silica bead DNA gel extraction kit (#K0513, ThermoFisher Scientific). The benzyldimethylalkylammonium chloride (BAC) method was used for native spreading of the DNA on a water surface and then loading it on carbon-coated 400-mesh magnetic nickel grids. After the spreading procedure, the electron density of the DNA was increased by platinum coating with the platinum-carbon rotary shadowing technique using the MED 020 High Vacuum Evaporator (Bal-Tec). The grids were then scanned in a semi-automated fashion using a transmission electron microscope (FEI Thalos 120, LaB6 filament) at high tension ≤ 120 kV and pictures were acquired with a bottom mounted CMOS camera BM-Ceta (4000 × 4000 pixels). The images were processed with MAPS Version 3.14 (ThermoFisher Scientific) and analyzed using MAPS Offline Viewer Version 3.14.11 (ThermoFisher Scientific). Mean±SD values of analyzed frequency of RIs obtained from three independent replicates were plotted and statistics was performed with Prism using unpaired *t*-test. The percentage of RFs containing ssDNA stretches was evaluated by manual scoring. Samples from mock-treated and HU-treated cells were pooled together for the analysis of ssDNA-containing RFs.

## QUANTIFICATION AND STATISTICAL ANALYSIS

Statistical parameters including sample size, number of biological replicates, applied statistical tests and statistical significance are reported in the corresponding figure legends or materials and methods section.

## AUTHOR CONTRIBUTIONS

D.D., M.L. and S.R. conceived the project. D.D., M.L., F.V., E.G., H.D, M.G.F., A.D., J.A.S., I.K., M.S., R.B., N.A.M. performed and analyzed experiments. S.T.D. and J.V.F. performed the CRISPR screen in Fig. 4b. M.D. performed the Cisplatin resistance genetic screen in Fig. 1c. A.N.K. generated the U-2OS-Tet^ON^GFP-CtIP^WT^ and U-2OS-Tet^ON^GFP-CtIP^8A^ cell lines. C.S.S., M.L., A.A.S, J.J. and S.R. were involved in supervision and funding acquisition. D.D. and S.R. wrote the manuscript. All the authors revised the final manuscript.

## ACKNOWLEDGEMENTS

We wish to thank Hana Hanzlikova and Lea Lingg for critical reading of the manuscript. We thank all the members of the Rottenberg lab for helpful discussion. We thank Horizon Discovery for performing and analysing the CRISPR-Cas9 screens in SW620, HT29 and BT549 cells. We thank Daniel Durocher for sharing RPE1-h*TERT TP53*^*-/-*^ and RPE1-h*TERT TP53*^*-/-*^*;BRCA1*^*-/-*^ cells. We are grateful to Georgina Hayoz and Fabiana Steck for their support at the Vetsuisse Faculty mouse facility and Denise Howald for her technical assistance. Financial support came from the Swiss National Science Foundation (310030_179360 to S.R., 31BL30_189698), 310030_208143 to A.A.S., the European Union (ERC-2019-AdG-883877 to S.R.), the Swiss Cancer League (KFS-5519-02-2022 to S.R.), the Boehringer Ingelheim Fonds (PhD fellowship to M.L.), and the Wilhelm Sander Foundation (no. 2019.069.1 to S.R.). Figures were created with Biorender.

## EXTENDED DATA FIGURE LEGENDS

**Extended data Fig. 1. a**, *H2afx* allelic modification rate in KB2P3.4 cells evaluated by TIDE analysis. **b**, H2AX immunoblotting in KB2P3.4-depleted cells. **c**, *H2afx* allelic modification rate in KB1P-G3 cells evaluated by TIDE analysis. **d**, H2AX immunoblotting in KB1P-G3-depleted cells. **e**, *H2afx* allelic modification rate in a competition assay in KB2P1.21 cells evaluated by TIDE analysis. **f**, Clonogenic survival assay of RPE1-h*TERT TP53*^*-/-*^*;BRCA1*^*-/-*^-derived cells treated, or mock treated, with the indicated concentrations of the PARPi olaparib for 12 days. **g**, H2AX immunoblotting in KB1P-G3 after genetic complementation. **h**, Clonogenic survival assay of KB2P3.4-derived cells treated, or mock treated, with the indicated concentrations of cisplatin for 12 days. Plotted values are the mean±SD clonogenic survival (n=3). *P*-values were calculated with the unpaired two-tailed Student’s *t*-test, ^****^ p<0.001. **i**, Clonogenic survival assay of KB2P3.4-derived cells treated or not with the indicated IR doses and stained after 12 days.

**Extended data Fig. 2. a**, KB1P-G3-derived cells were treated with olaparib (3μM) for 24h. Plotted values are the mean±SD of micronucleated cells (n=2). *P*-values were calculated with the unpaired two-tailed Student’s *t*-test. **b**, DNA fiber analysis in KB2P3.4-derived cells treated according to the depicted scheme. Plotted values show the median of individual IdU/CldU ratios from at least 250 fibers (n=3). ^****^ P<0.0001 unpaired One-way Anova test. **c**, Cell growth assay in KB2P3.4-derived cells. **d**, DNA fiber analysis in KB2P3.4-derived cells treated according to the depicted scheme. Plotted values show the median of individual IdU/CldU ratios from at least 170 fibers (n=3). ^****^ p<0.001 unpaired One-way Anova test. **e**, MRE11 SIRF in KB2P3.4 cells treated with HU as in Fig. 3c. Plotted values are the number of MRE11 SIRF foci/cell and the median from at least 300 cells (n=3). *P*-values were calculated with the unpaired two-tailed Student’s *t*-test.

**Extended data Fig. 3. a**, Clonogenic survival assay of KB2P3.4-derived cells treated, or mock treated, with the indicated concentrations of the PARPi olaparib for 12 days with and without the indicated concentrations of the ATMi AZD0156. Plotted values show the mean±SD of clonogenic survival (n=3). **b**, CtIP protein map showing all the validated and predicted ATM phosphorylation sites. **c**, Immunoblot showing GFP-CtIP^WT^ or GFP-CtIP^8A^ expression in U-2OS-TeT^ON^-CtIP cells.

