## Supplementary figures and images for "H2AX promotes replication fork degradation and chemosensitivity in BRCA-deficient tumours"

### Supplementary Figure 1

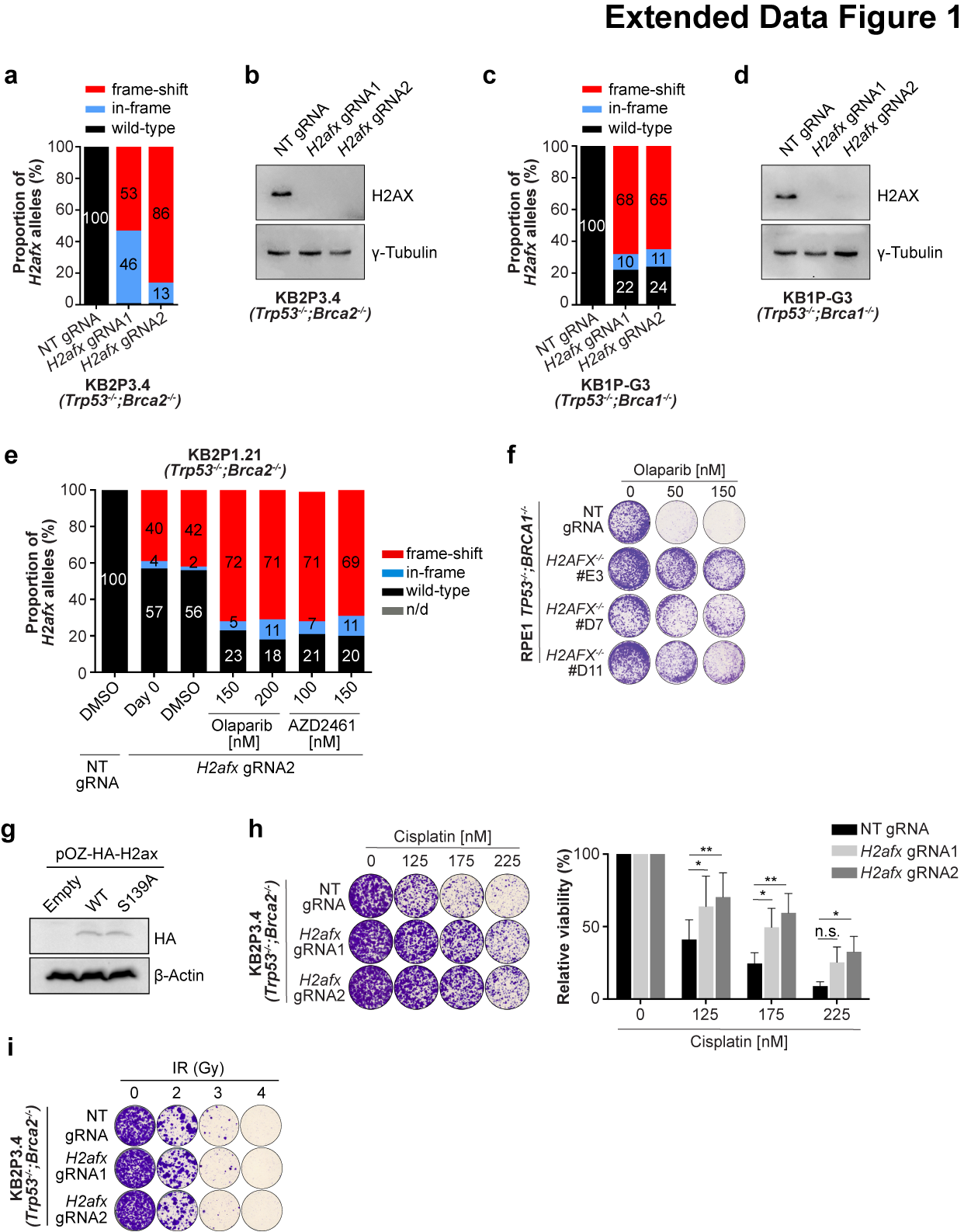

### Supplementary Figure 2

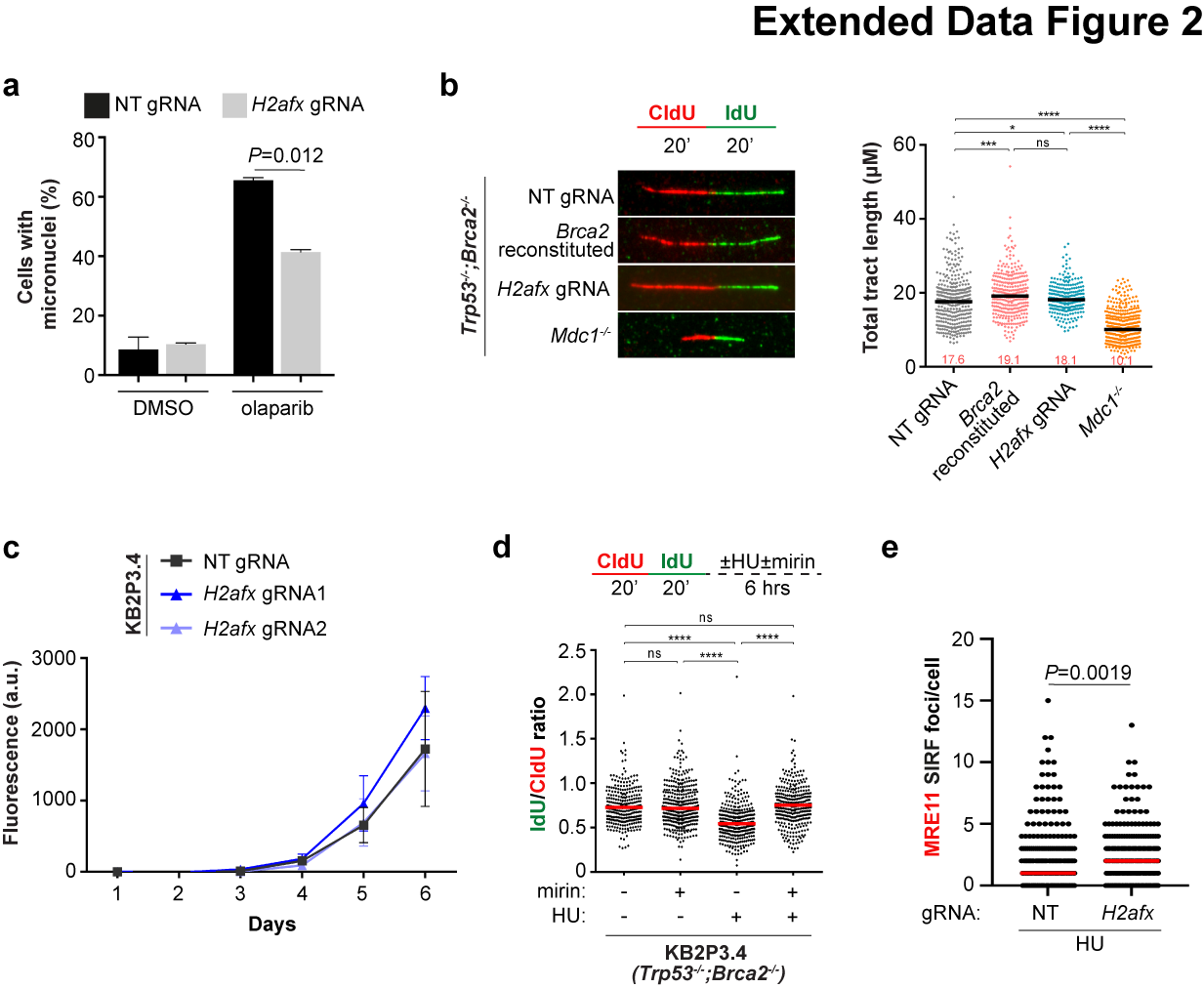

### Supplementary Figure 3

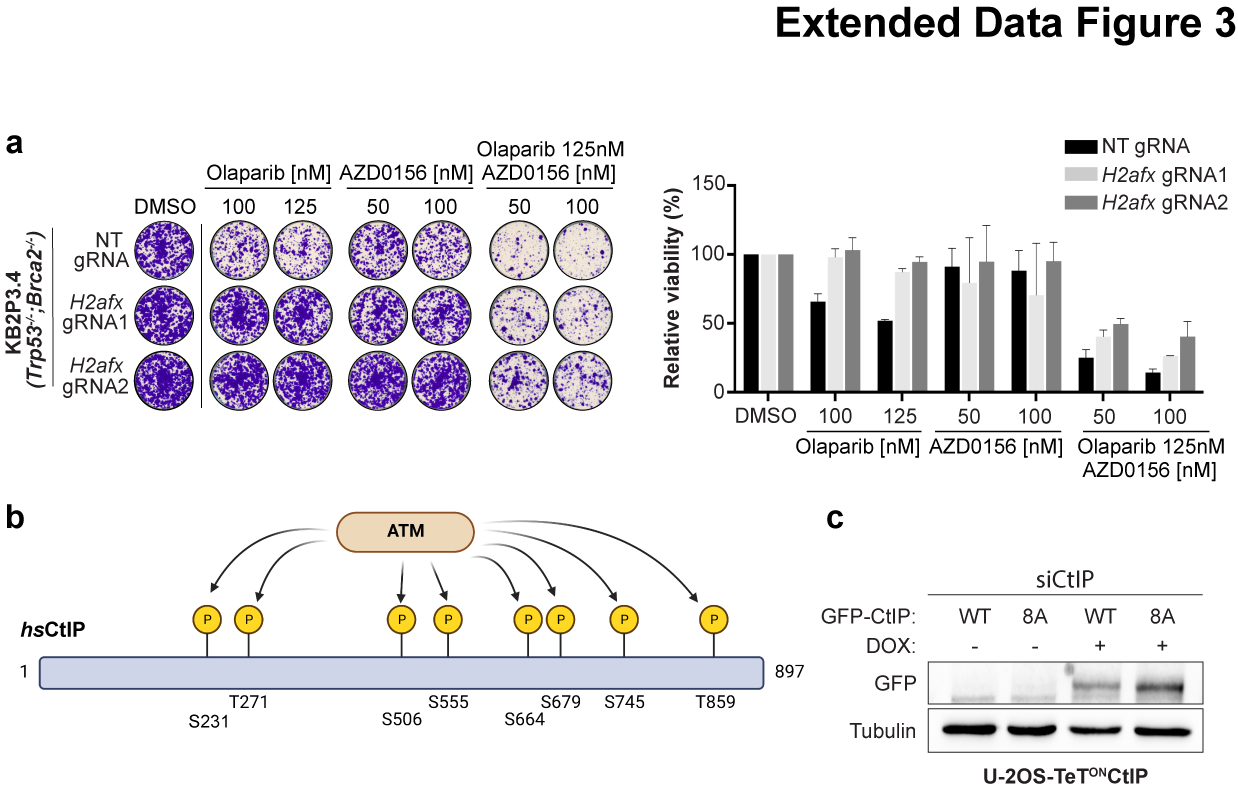
